# Monoclonal antibodies against cell wall chitooligomers as accessory tools for the control of cryptococcosis

**DOI:** 10.1101/2021.06.11.448165

**Authors:** Alexandre Bezerra Conde Figueiredo, Fernanda L. Fonseca, Fernando de Paiva Conte, Marcia Arissawa, Marcio L. Rodrigues

## Abstract

Therapeutic strategies against systemic mycoses can involve antifungal resistance and significant toxicity. Thus, novel therapeutic approaches to fight fungal infections are urgent. Monoclonal antibodies (mAbs) are promising tools to fight systemic mycoses. In this study, mAbs of the IgM isotype were developed against chitin oligomers. Chitooligomers derive from chitin, an essential component of the fungal cell wall and a promising therapeutic target, as it is not synthesized by humans or animals. Surface plasmon resonance (SPR) assays and cell-binding tests showed that the mAbs recognizing chitooligomers have high affinity and specificity for the chitin derivatives. *In vitro* tests showed that the chitooligomer mAbs increased the fungicidal capacity of amphotericin B against *Cryptococcus neoformans*. The chitooligomer-binding mAbs interfered with two essential properties related to cryptococcal pathogenesis: biofilm formation and melanin production. In a murine model of *C. neoformans* infection, the combined administration of the chitooligomer-binding mAb and subinhibitory doses of amphotericin B promoted disease control. The data obtained in this study support the hypothesis that chitooligomer antibodies are of great potential as accessory tools in the control of cryptococcosis.

## Introduction

Fungal infections affect more than one billion people, resulting in approximately 13.5 million life-threatening infections and more than 1.7 million deaths annually (1). Aproximately 30% of patients diagnosed with histoplasmosis living with HIV/AIDS die in Latin America (2). Each year 220,000 new cases of cryptococcal meningitis occur worldwide, resulting in approximately 180,000 deaths (3). More than 400,000 cases of *Pneumocystis* pneumonia occur in Africa each year, with an estimated fatality rate of 100% if the disease is not treated (3). Although these numbers are alarming, there are only a few options for the treatment of fungal diseases. Immunotherapeutic strategies have been proposed as tools to fight fungal diseases. Antibodies with therapeutic potential were developed against *H. capsulatum* histone 2B (4), melanin (5), and heat shock proteins (6); *C. albicans* and *A. fumigatus* β-glucans (7); *C. neoformans* glycosylceramide (8), melanin (9), and glucuronoxylomannan (10), among others.

Chitin is essential for the integrity of fungal cell walls (11). Since this polysaccharide is not synthesized by humans or animals, chitin is a promising candidate for the antifungal therapy (11). The inhibition of chitin synthesis in fungi is not trivial, due to the general redundancy of genes regulating chitin formation in fungal cells (12). Chitin oligomers or chitooligomers are formed by the partial enzymatic hydrolysis of chitin in fungal cells (13). In *C. neoformans*, the blocking of chitooligomers with a lectin prior to injection of the fungus in mice reduced brain colonization (14), suggesting that these structures could be targeted by antifungal approaches. Monoclonal antibodies (mAbs) to chitooligomers are still not available, which limits the evaluation of these molecules as therapeutic tools.

In this study, mAbs were developed against chitooligomers using the hybridoma technique. These mAbs recognized chitooligomers on the surface of *C. neoformans* and *Candida albicans*. Surface plasmon resonance (SPR) assays demonstrated that the mAbs are avid and specific for the chitooligomers. *In vitro*, these mAbs inhibited fungal growth when combined with amphotericin B. Moreover, one the chitooligomer-binding mAbs was highly effective to control cryptococcosis in a mice model of infection, in the presence of subinhibitory amphotericin B. Our findings suggest that chitooligomer-binding mAbs are promising as accessory tools to fight cryptococcosis.

## Results

### Generation of mAbs against chitooligomers

For immunization of mice, we first challenged the animals with *C. deuterogattii* followed by 2 intraperitoneal injections at 15-day intervals with the β-1,4-linked N-acetylglucosamine trimer, chitotriose, in aluminum hydroxide. Mice were then challenged intravenously with chitotriose in PBS 3 days after the last intraperitoneal injection. Fifity eight hybridomas reacting with chitotriose were selected, and finally the 10 hybridomas that showed the highest response were selected for ELISA using chitotriose-BSA as the primary antigen. The most reactive antibodies, accordingly to the ELISA results, were purified and sequenced (Table 1), and 2 mAbs (namely DD11 and CC5) were identified with different complementarity determining regions (CDR), which were characterized as of the IgM subtype (Figure 1). The light and heavy regions (VL and VH) were characterized with bands of approximately 23 kda and 70 Kda, respectively (Figure 1).

**Table 1:** CDR sequence comparisons of the chitooligomer mAbs

| mAb | CDR1 | CDR2 | CDR3 |
| --- | --- | --- | --- |
|  | Variable light chain (VL) |  |  |
| **CC5** | *SASSSISYMH* | *DTSKLAS* | *QRSSYPCT* |
| **DD11** | *SASSSVSYMH* | *STSNLAS* | *QQRSSYPLT* |
|  | Variable heavy chain (VH) |  |  |
| **CC5** | *GFTFSDAWMD* | *IRSKANNHA* | *HRYDGFDY* |
| **DD11** | *GFTFSDYGMA* | *ISNLAYSIY* | *DYYGSSYWYFDV* |

**Figure 1:**
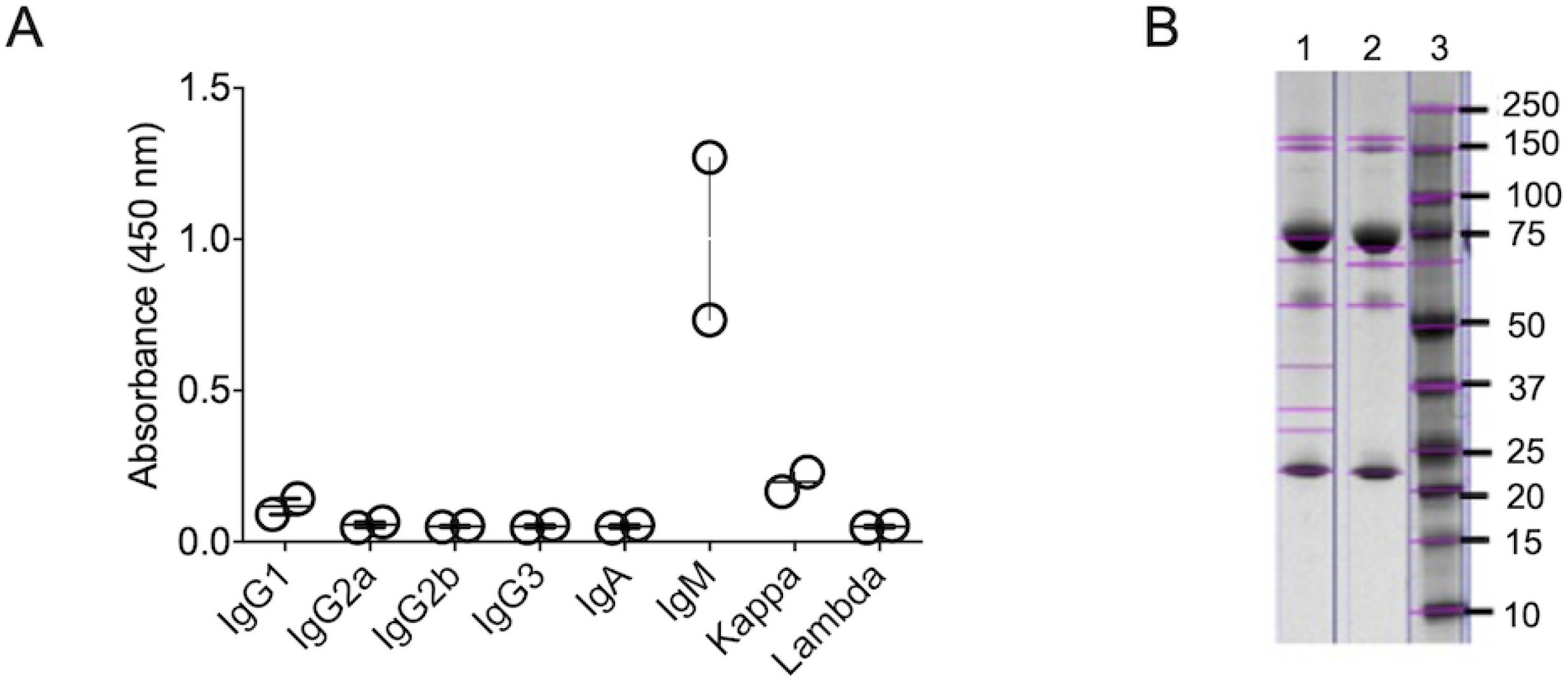
Characterization of mAbs to chitooligomers. (A). Isotyping of mAbs DD11 and CC5 was performed using culture supernatants of the hybridomas producing each antibody. Results from each hybridoma were compiled and shown in panel A, where each sphere represents the signals obtained from each antibody. The highest signals were obtained for IgM, with mAb DD11 producing the highest reacitivty. (B) Denaturing gel of purified mAbs denoting the predominant bands corresponding to the heavy (∼ 70 kDa) and light (∼ 23-24 kDa) chains of IgM. Lane 1 shows the electrophoretic separation of mAb CC5, while lane 2 represents mAb DD11. Line 3 represents the molecular mass markers.

### Antigen recognition assays

The mAbs were immobilized by amine coupling chemistry, and chitotriose was tested at two concentrations (0.1 and 0.06 mM) to analyze the association (kA) and dissociation constants (kD). SPR was adjusted to a 1:1 interaction model and, finally, kA and kD were determined for each antibody. Regardless of the concentration of the antigen, mAb DD11 demonstrated higher affinity than CC5 (Table 2). Both mAbs were tested against other molecules (glycine and BSA), and they showed no affinity or specificity for these molecules (Figure 2). We also included cell-binding assays in our mAb characterization. In these assays, *C. neoformans* and *C. albicans* were tested through an adaptation of conventional ELISA to allow the use of intact cells (Figure 3A-B). The mAbs gave positive reactions above a cut off of 10^3^ cells/mL for both fungi. The reactions were considered as positive when they produced spectrophotometric values 3-fold greater than the negative control. The mAbs were also tested for their ability to recognize other cell types (Figure 3C-D), including human pulmonary cells (A549), Gram-positive bacteria (*Staphylococcus aureus*), Gram-negative bacilli (*Escherichia Coli)*, and parasites (*Giardia lamblia)*. The mAbs did not react with *G. lamblia*, A549 cells, *E. coli* or *S. aureus* at 10^4^ cells/mL. We also used dot blot assays to test the mAb-fungi interactions. In these tests, the mAbs required at least 10^6^ cells/mL to recognize *C. neoformans*. However, the DD11 mAb required only 10^4^ cells/mL to recognize *C. albicans*, in contrast to CC5 (Figure 3E-F). Once again, reactions were considered to be positive when values were 3-fold greater than the control cut-off.

**Table 2:**
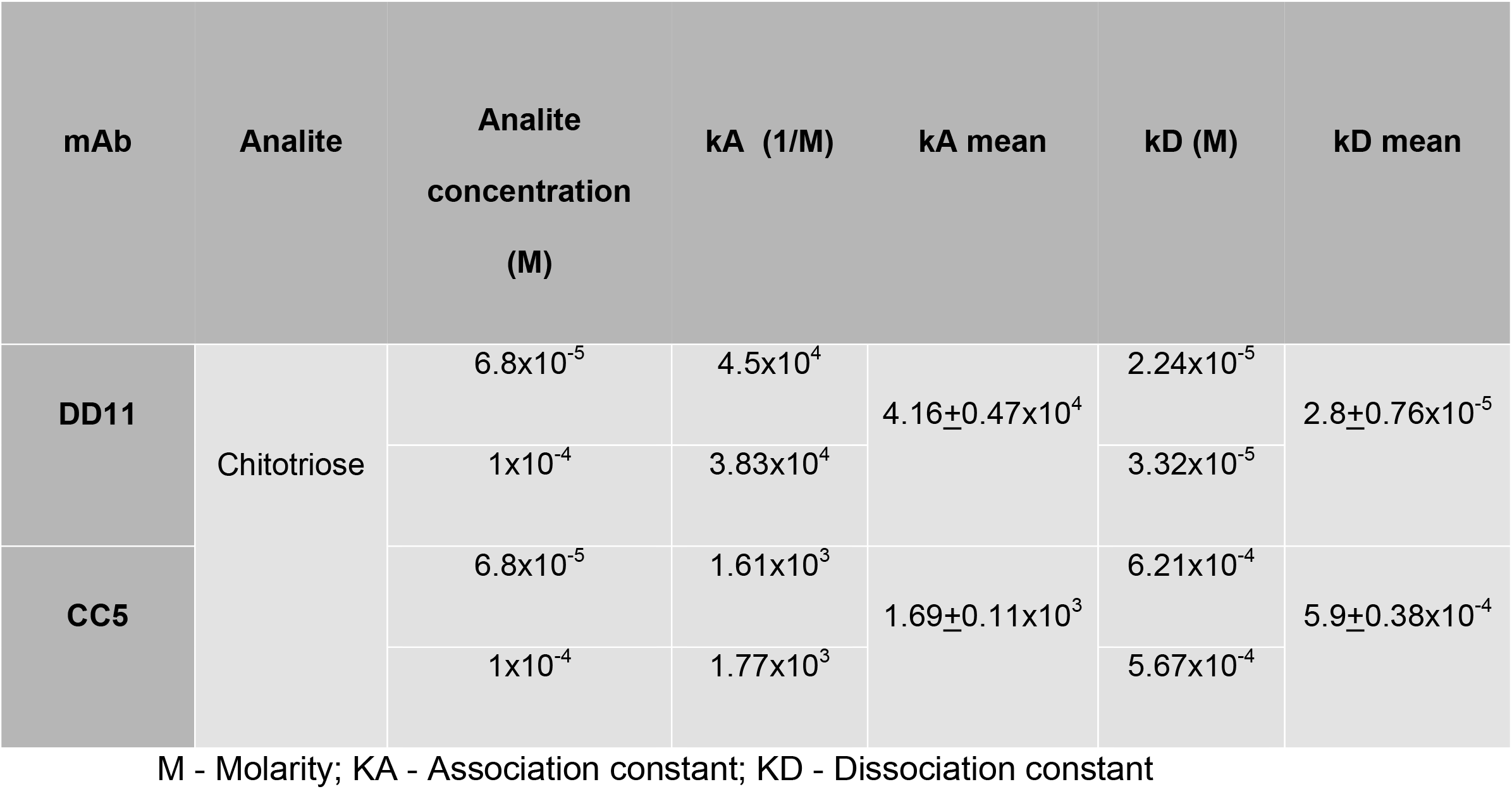
Kinetics of the binding of chitooligomer mAbs to chitotriose

**Figure 2:**
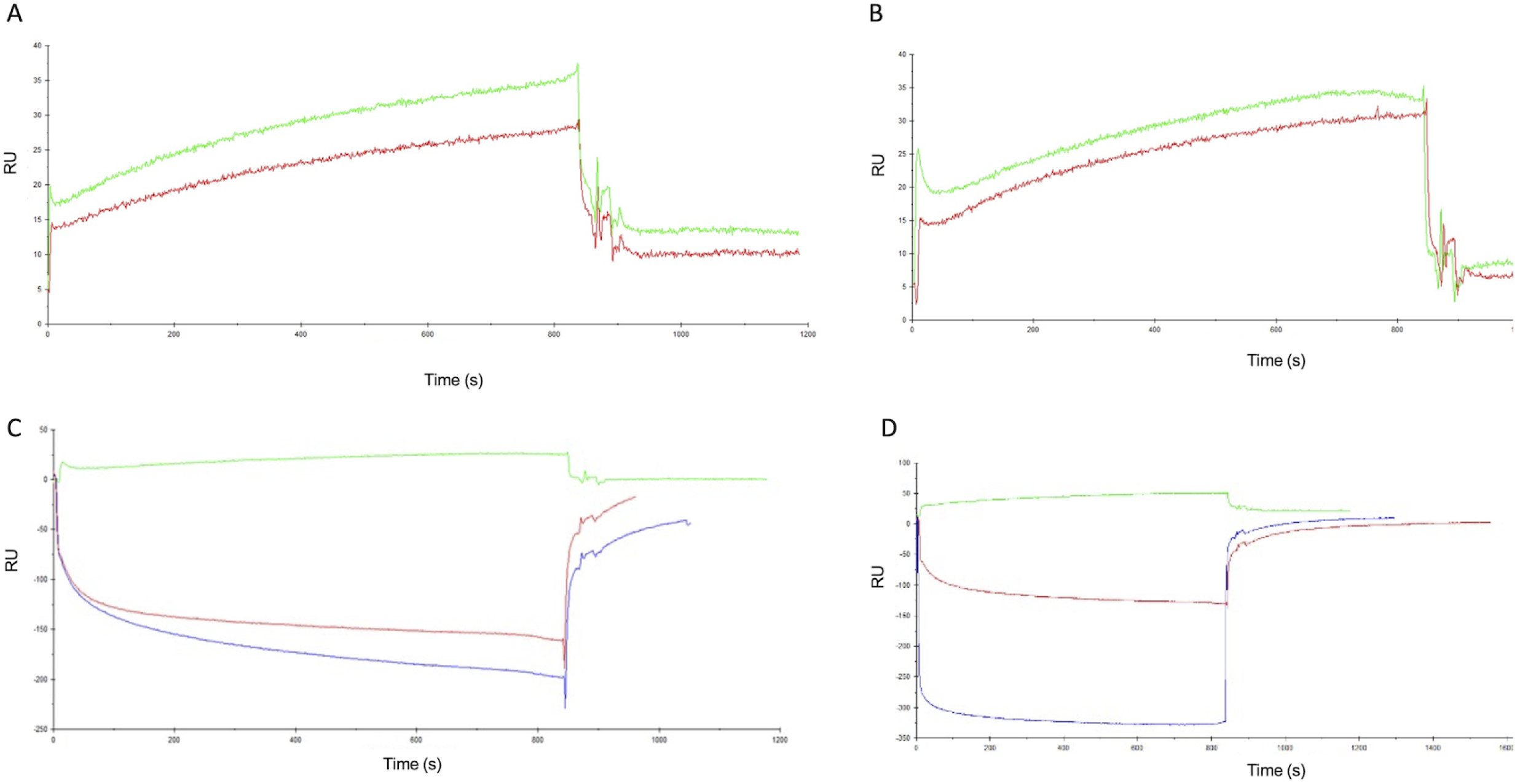
SPR sonogram representative of the interaction of the chitooligomer mAbs with chitotriose. The ligands tested were the mAbs DD11 (A) and CC5 (B). C and D represent the negative controls for mAbs DD11 and CC5, respectively. The surfaces of the flow cells were activated and the ligands immobilized at 100 μg/mL in 10 mM sodium acetate, pH 5.0. In panels A and B, red and green lines correspond to the interaction of the mAbs with of 0.06 nM and 0.1 nM chitotriose, respectively. In panels C and D, the green lines represents chitotriose (0.06 nM), while red and blue lines represents BSA and glycine in the same concentration, respectively. Panels C and D represent the controls of each antibody (C, mAB DD11; D, maB CC5). The analyte injection corresponds to time zero. The rise of the curves represent analyte-substrate binding. Approximately 800 seconds after injection, disassociation starts, resulting in the abrupt drop of the curves. RU represents response units generated by the equipment’s software.

**Figure 3:**
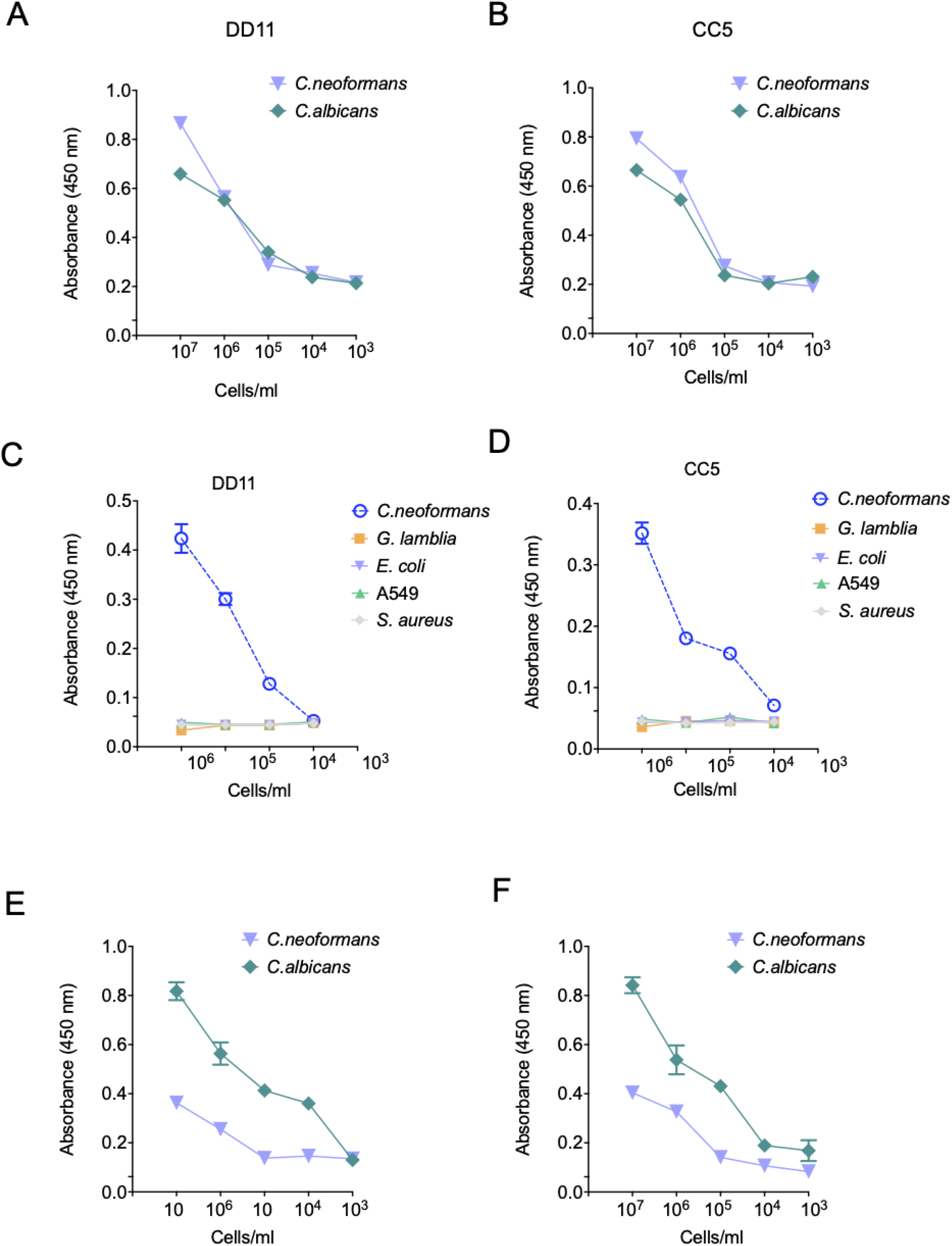
Binding of chitooligomer mAbs to whole cells of *C. neoformans* and *C. albicans*. The wells of ELISA plates were coated with cells of *C. albicans* (blue lines) or *C. neoformans* (green lines) at cell densities ranging from 10^2^ to 10^7^/mL. The reactivity of mAbs DD11 (A) and CC5 (B) at of 12.5 µg/mL are shown. Similar tests were performed in C (mAb DD11) and D (mAb CC5) with *G. lamblia* (green lines), A549 human cells (orange lines), Gram-negative *E. coli* (purple lines) and Gram-positive *S. aureus* (grey lines). Dot blot also the binding of chitooligomer mAbs to whole cells of *C. neoformans* and *C. albicans*. MAbs DD11 (E) and CC5 (F) were tested against whole cells of *C. albicans* (blue lines) and *C. neoformans* (green lines) at varying cell densities. The mAbs were used at 12.5 µg/mL. The results illustrate a representative experiment of three independent replicates producing similar results.

### Effects of the chitooligomer mAbs on the formation of cryptococcal biofilms

Due to the roles of fungal biofilms in pathogenesis (15), we tested the effect of the chitooligomer mAbs on the formation of *C. neoformans* biofilms. At 12,5 μg/mL, the mAbs significantly inhibited the formation of biofilms, and also affected mature biofilms (p <0.05; Figure 4). As a control of biofilm inhibition, amphotericin B was used at 1 μg/mL. Lower mAb concentrations were also tested; however, no effects were observed. Similar results were obtained for *C. albicans* biofilms (data not shown).

**Figure 4:**
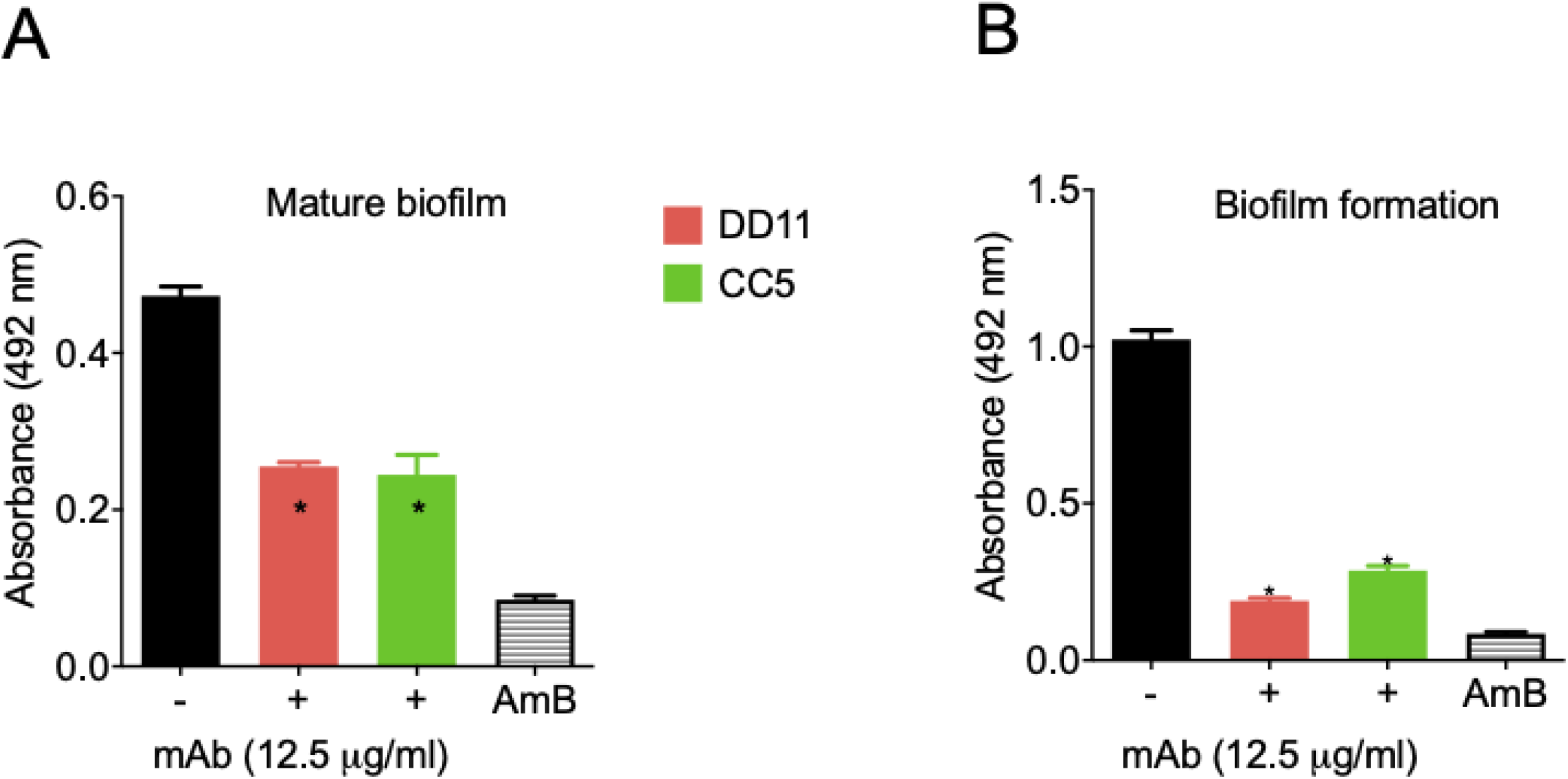
Effect of mAbs to chitooligomers on cryptococcal biofilms. Mature biofilm and biofilm formation had their metabolic activity measured indirectly by the XTT reduction assay. MAbs DD11 (red), CC5 (green), and amphotericin B (AmB, 1 µg/mL) were tested for their ability to affect biofilms. A illustrate the treatment of mature biofilms with mAbs and B illustrate the incubation of *C. neformans* with the mAbs before biofilm formation. Data illustrate a representative experiment of three independent replicates producing similar results. Asterisks denote statistically significant differences (p < 0.01) in comparison to control systems.

### Chitooligomer mAbs affect the melanization of *C. neoformans*

The capacity of *C. neoformans* to produce melanin represents an important virulence factor (16). Considering that the mAbs tested here target components of the cell wall, where melanin is deposited, we evaluated the ability of these antibodies to affect pigmentation. MAbs DD11 and CC5 were tested at concentrations ranging from 0.2 to 25 μg/mL, and L-DOPA was used as a substrate for melanization. The DD11 mAb partially inhibited cryptococcal melanization at concentrations higher than or equal to 6.2 μg/mL (p <0.05), while at lower concentrations no inhibition was detected (p> 0.05). At concentrations greater than 6.2 μg/mL (p <0.001), the CC5 mAb also prevented melanization completely, while at 3.2 μg/mL a partial inhibition occurred. Lower concentrations had no effect on the fungal pigmentation (p> 0.05, Figure 5).

**Figure 5:**
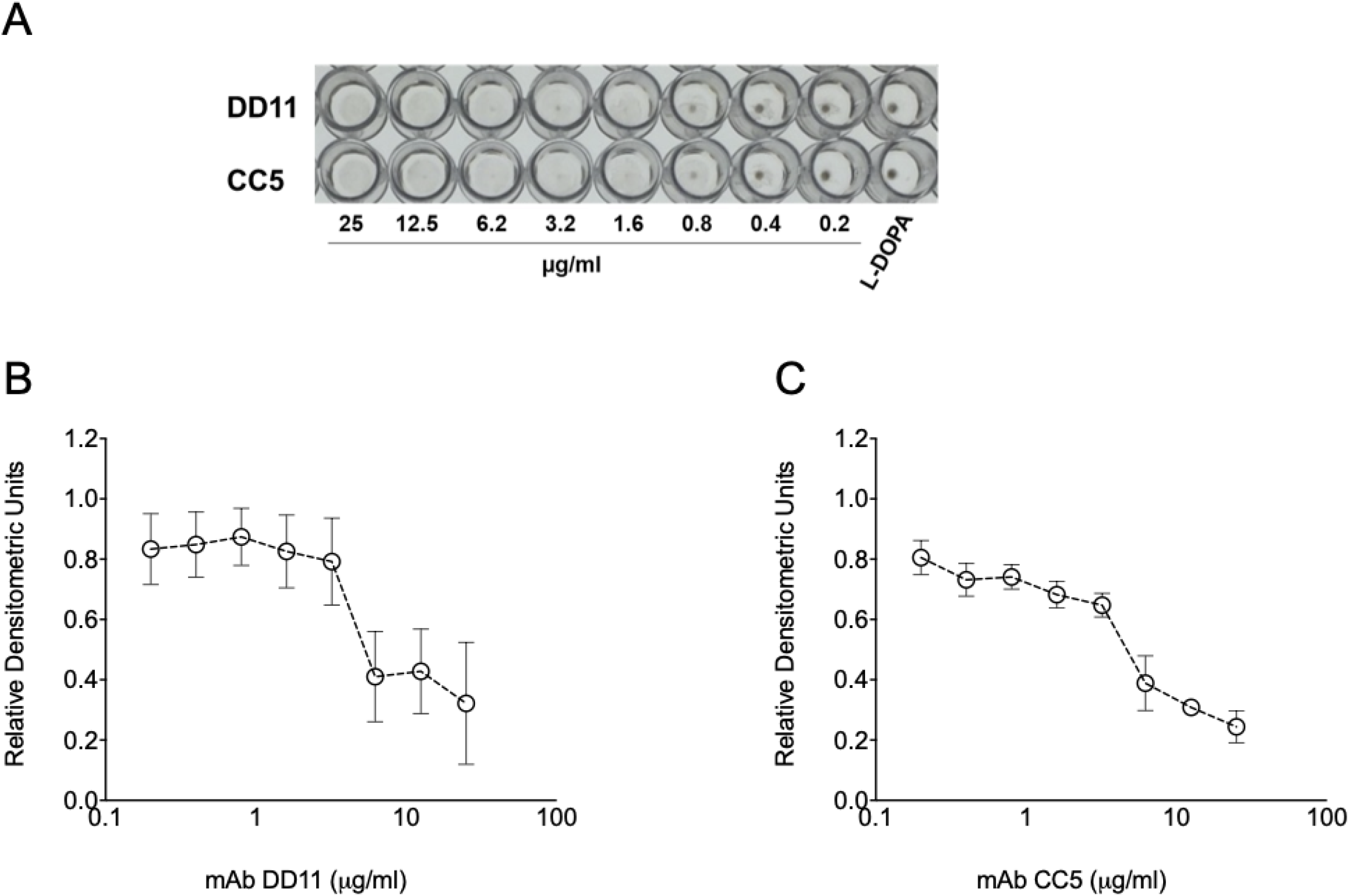
Effect of the chitooligomer mAbs on the melanization of *C. neoformans*: (A) Pigmentation of *C. neoformans* in presence of mAbs DD11 and CC5. Pigmentation in *C. neoformans* was visually assessed by sedimentation of brown to black color at the bottom of the 96-well plates. B and C illustrate the densitometric quantification of pigments treated with mAbs DD11 and CC5, respectively. The results illustrate a representative experiment of three independent replicates producing similar results.

### Antifungal activity

The potential antifungal activity of the mAbs against *C. neoformans* was tested using standardized methods proposed by the European Committee on Antimicrobial Susceptibility Testing (EUCAST) (17). When tested alone, the mAbs did not show any effect on the fungal growth (Figure 6). We then tested the mAbs in the presence of subinhibitory (0.1 μg/mL) amphotericin B. In the presence of the antifungal, a dose-dependent inhibition of fungal growth was observed for both antibodies. The effects of the mAbs and amphotericin B were synergistic (Table 3). We also observed an additive effect of amphotericin B with mAb DD11 at 1.6 μg/mL. These calculations were based on the fractional inhibitory index (FII).

**Table 3.** Antifungal effects of chitooligomer mAbs in combination with amphotericin B*.

| mAb | Amphotericin B<br>[ 0.1 µg/mL] | IIF |
| --- | --- | --- |
| DD11 [µg/mL] | 12.5 | 0.8 |
|  | 6.2 | 0.7 |
| CC5 [µg/mL] | 12.5 | 0.8 |
|  | 6.2 | 0.9 |

\*Additive or synergistic effects were determined on the basis of the calculation of the fractional inhibitory index. Values corresponding to 1.0 represent an additive effect, while values corresponding to <1.0 denote a synergistic effect (46).

**Figure 6:**
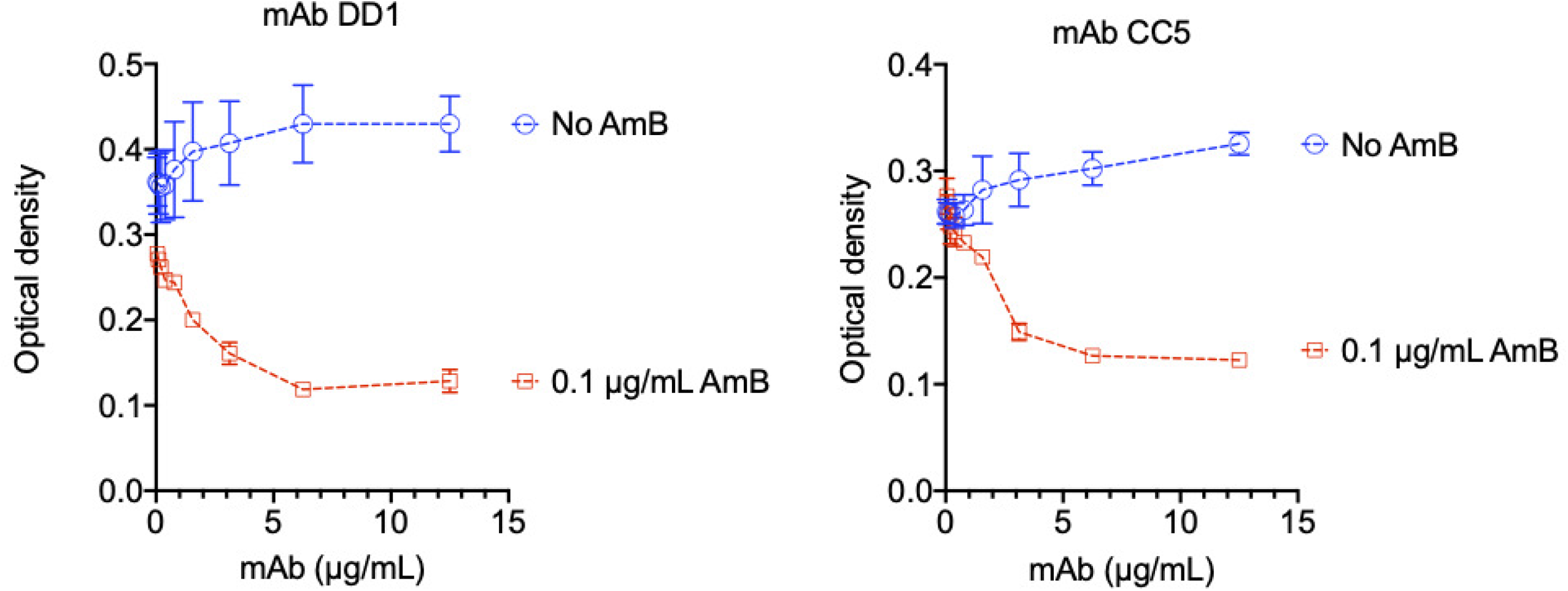
Analysis of cryptococcal growth in the presence of subinhibitory amphotericin B and mAbs DD11 or CC5. *C. neoformans* cells were incubated in the absence or presence of mAbs, both at concentrations varying from 0 to 12.5 μg/mL. Alternatively, fungal cells were cultivated in the presence of subinhibitory amphotericin B (AmB 0.1 μg/mL) alone (no mAb, 0 μg/mL) or in combination with the mAbs. The combination of the mAbs with subinhibitory amphotericin B resulted in inhibition of fungal growth. The results illustrate a representative experiment of three independent replicates producing similar results. Asterisks denote statistically significant differences.

### The association of a chitooligomer mAb and subinhibitory amphotericin B results in the control of animal cryptococcosis

On the basis of the antifungal effects of the association between the chitooligomer mAbs and subinhibitory amphotericin B, we tested the ability of this association to control mice cryptococcosis (Figure 7). MAb DD11 was selected for these assays due to its broader activity (synergistic and additive effects) in the presence of amphotericin B. The mortality rate of untreated animals infected with *C. neoformans* (control system) was 100% on the 28th day after infection. Similarly, the treatment of lethally infected mice with the mAb alone at different concentrations resulted in 100% death at day 29 post infection. All animals placed in the group treated only with amphotericin B (0.25 mg/kg) died at day 37 post infection, while all animals placed in the group treated with a combination of mAb DD11 (85 μg/animal) with amphotericin B (0.25 mg/kg) survived until the experiment was interrupted at day 90 post infection. In the control group treated with the standard dose of amphotericin B (2.5 mg / kg) no animals died (data not shown).

**Figure 7:**
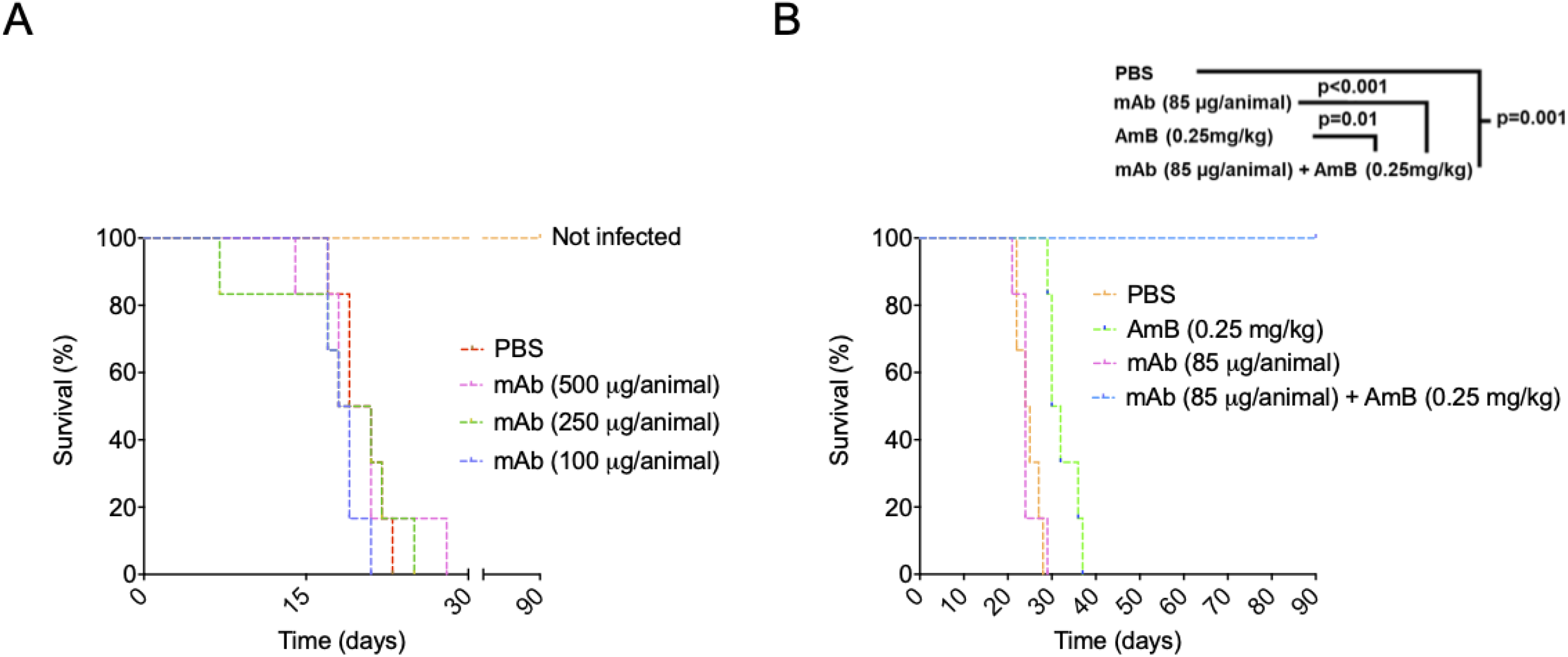
An association of mAb DD11 with subinhibitory amphotericin B (AmB) improves the survival of lethaly infected mice in a mouse *C. neoformans* model. A. Balb/c mice (n=7) receiving PBS or mAb DD11 died at days 22 to 29 post infection with *C. neoformans*. B. Animals receiving subinhibitory AmB (0.25 mg/kg) post infection and died at day 37 post infection. All mice receiving the combination of subinhibitory AmB and the mAb survived until the experiment was interrupted (p = 0.001).

## Discussion

Several studies have shown that monoclonal antibodies can be protective against fungal infections (7). Proteins, polysaccharides, pigments, and even fungal glycolipids were characterized as the targets of protective antibodies (6–8, 18–21). Our laboratory previously demonstrated that blocking cell wall chitooligomers is beneficial for the control of animal cryptococcosis (22), but mAbs raised to chitin or its derivatives are still not available. Thus, we chose chitotriose, a water-soluble chitin oligomer composed of three units of β1,4-linked N-acetyglucosamine, as a target for developing mAbs.

The immunization strategy using *C. deuterogattii* whole cells was adopted based on our previous study showing that chitooligomers are exposed on the outer layers of the cryptococcal surface (23). This immunization strategy, which certainly stimulated immune responses to multiple antigens, was followed by sequencial boosts of the humoral response to the chitooligomers using chitotriose combined with AlOH_3_ (24, 25). In contrast to IgM, IgG was not detected, suggesting a greater proliferation of memory cells of the IgM type (26, 27). We assumed that their rapid expansion followed by the generation of plasma cells independent of T cells led to a greater accumulation of IgM-producing cells, probably due to the stimulation caused by chitotriose (28).

Amphotericin B, the main drug to fight cryptococcosis and other systemic mycoses, is toxic and expensive (29). Therefore, therapeutic protocols using smaller doses of amphotericin B have the potential to be more affordable and produce less accentuated collateral effects. The chitooligomer mAbs had no effects on the fungal growth alone. However, cryptococci were more susceptible in vitro to amphotericin B in the presence of the chitooligomer mAbs. We still do not know why the antibodies render *C. neoformans* more susceptible to amphotericin B, but it is likely that the antibodies interfere with the cell wall, as suggested by their ability to affect melanization. Considering that amphotericin B acts on the plasma membrane (30), we speculate that changes in the cell wall could make the *C. neoformans* more permeable to the antifungal, increasing its efficacy. Interference with the cell wall could explain the effects on biofilm formation, considering the essential role of the fungal surface in this process (31).

The development of mAbs against fungal targets has been explored over the past two decades (18, 32, 33). Two examples are the mAbs 18B7 against *Cryptococcus* GXM and 2G8 against *Candida* β 1,3 glucan (34, 35). These antibodies vary in ther protective mechanisms: mAb 18B7 is an immunomodulator with no antifungal effects, while mAb 2G8 interferes with fungal growth (34, 35). Other mAbs raised to fungal components can promote growth inhibition (36), inhibition of biofilm formation (37), reduced adherence and germination (38), and inhibition of melanin formation (9). Our results and these reports suggest that the chitooligomer antibodies could facilitate infection control in mice without affecting the host’s physiology, as inferred from their affinity and specifity for fungal cells.

Considering the ability of the chitooligomer mAbs to participate in antifungal activity, biofilm formation and pigmentation, we asked whether these antibodies would be effective *in vivo*. In fact, mAb DD11 promoted the control of animal cryptococcosis in the presence of a subinhibitory concentration of amphotericin B. This combination resulted in 100% survival of animals infected with lethal inocula of *C. neoformans*. The mechanisms explaining these effects are still unclear, but they could be related to the effects of the antibody *in vitro*, in addition to the modulation of the immune response in favor of the host (39), and interference with the immunological recognition of chitooligomers and chitin. In this sense, we have previously demonstrated that blocking cryptococcal chitooligosaccharides by other approaches resulted in attenuated pathogenesis (14).

Our results support the notion that mAbs against chitooligomers might be useful as biopharmaceuticals for treating fungal diseases along with lower doses of amphotericin B. Thus, the mAbs developed in this study could be used as auxiliary tools in the therapy against fungal infections. Future studies with humanized antibodies on optimal doses and immunomodulation *in vivo* could be the basis for the formulation of more efficient protocols to fight cryptococcosis.

## Acknowledgements

M.L.R. is currently on leave from the position of associate professor at the Microbiology Institute of the Federal University of Rio de Janeiro, Brazil. M.L.R. is supported by grants from the Brazilian Ministry of Health (grant 440015/2018-9), Conselho Nacional de Desenvolvimento Científico e Tecnológico (CNPq; grants 405520/2018-2 and 301304/2017-3), and Fiocruz (grants PROEP-ICC 442186/2019-3, VPPCB-007-FIO-18, and VPPIS-001-FIO18). This study was financed in part by scholarships from the Coordenação de Aperfeiçoamento de Pessoal de Nível Superior (CAPES, Brazil, Finance Code 001). M.L.R. also acknowledges support from the Instituto Nacional de Ciência e Tecnologia de Inovação em Doenças de Populações Negligenciadas (INCT-IDPN). The funders had no role in the decision to publish or preparation of the manuscript.

## Transparency declaration

Part of the data presented here is the subject of a pending patent application (Ref.: BR 10 2020 002165 6).

## Materials and Methods

### Animal use and ethics statemen

Female Balb/c mice (6-8 weeks old) were obtained from the animal facility of Fiocruz (CECAL). All experimental procedures were approved by the Oswaldo Cruz Foundation’s Ethical Committee on the Use of Animals (CEUA, protocol: LW-13/16).

### Eukaryotic and prokaryotic cells

Fungal strains used in this study were *C. neoformans* (H99), *C. deuterogattii* (R265), and *C. albicans* (ATCC 90028). The microorganisms were maintained in Sabouraud broth (*C. neofomans*) or liquid brain heart infusion (BHI, *C. albicans*). For in vitro assays, the cells were cultured in a minimal medium (15 mM glucose, 10 mM MgSO_4_, 29.4 mM KH_2_PO4, 13 mM glycine, 3 μM thiamine-HCl, pH 5.5) and kept under rotation for 24 h at 37 °C. The cells were obtained by centrifugation, washed in PBS, and counted in a Neubauer’s hematocytomer. Antibody especifity tests included *Giardia lamblia* (ATCC 30957), A549 human cells (ATCC CCL-185), *Escherichia coli* (ATCC 9637), and *Staphylococcus aureus* (ATCC 25923).

### Mice immunization

Mice (female Balb/C, 6 weeks old) were immunized intraperitoneally (i.p.) every 14 days according to Guimarães et al. with some adaptations (40). The animals were immunized via i.p. with *C. deuterogattii* (strain R265, 1 × 10^6^ cells/mL, 100 µl/animal) previously fixed with 4% paraformaldehyde (PFA) and washed in 10 mM phosphate saline (PBS), pH 7.4. This procedure was followed by two immunizations via i.p. with an interval of 14 days with 200 μg of chitotriose combined with aluminum hydroxide (1.5 mg, 100 µl/animal). Finally, the animals were immunized intravenously (i.v.) with 50 μg of free chitotriose 3 days after the last immunization, using PBS as a vehicle (50 µl/animal). Bleeding was performed at the end of the immunization to check the serum antibody titer by indirect ELISA. The pre-immune sera from all animals was used as controls, and reaction cut-off for the screenings. After the four immunizations, the animals were submitted to splenectomy and splenocyte processing to perform cell fusion with murine myeloid cells (SP2/0)(41). Splenocytes and SP2/0 cells at the proportion of 1:2 were fused in the presence of 50% (v/v) polyethylene glycol (PEG) 3000-3700, preheated to 37°C. Subsequently, the cell homogenate was suspended in DMEM containing glutamine (6.4 mM), 100 units/mL penicillin, 0.1 mg/mL streptomycin, and 0.25 μg/mL amphotericin B, and 20% fetal bovine serum (SFB). The cells were transferred to 96-well plates (100 μL/well). Three wells were reserved as controls of the selection medium (DMEM with hypoxanthine, aminopterin and thymidine; HAT), to which 6 × 10^4^ SP2 / 0 cells were added. The plates were incubated at 37°C, in an atmosphere of 5% CO_2_ for 24 h. After the initial incubation, 100 μL of the selection medium (DMEM / 6.4 mM Glutamine / 1x ATB / 20% SFB / 2x HAT) was added to the wells to select viable, hybrid cells. After 14 days, the culture supernatants were used in indirect ELISA using chitotriose as the primary antigen.

### ELISA for the determination of serum titers

ELISA (96-well) plates were coated with chitotriose complexed with serum bovine albumin (chitotriose-BSA, Vector Laboratories – Cat.:G-5000) at 0.5 μg/mL in PBS, and incubated overnight at 4°C. Subsequently, the plate was incubated with PBS containing 1% BSA for 1 h at 37°C followed by washing with PBS containing 0.05%. Tween. The sera of the animals were added at different dilutions and incubated for 2 h at 37°C. The plate was washed three times with PBS-Tween with subsequent addition of anti-murine IgG and anti-IgM antibodies conjugated to peroxidase, followed by incubation for 2 h at 37°C. The plates were then washed as described above, and incubated with tetramethylbenzidine (TMB) for 30 min at 37°C. The reaction was stopped with 1 M HCl, and the spectrophotometric readings were obtained at 450 nm. The reactions considered as positive were those with absorbance values corresponding to 3 times the cut-off, without the addition of primary antibodies. For determining polyclonal and monoclonal antibody reactivity, the same procedure was performed, with the difference that the primary antibody of the reaction came from the hybridoma culture supernatants.

### Cloning of polyclonal hybridomas

The cloning of positive polyclonal hybridomas in the ELISA assay was performed by limiting dilution (20). The cells were incubated at 37°C, in a 5% CO_2_ atmosphere for 14 days, and clonality (monoclonal or polyclonal) was observed from the 5^th^ day. The cultures that remained viable had their supernatants tested by ELISA using chitotriose-BSA as the primary antigen. Once again, the reactions were considered as positive when the absorbance values corresponded to 3 times the cut-off, without the addition of primary antibodies. The isotyping of the selected clones was performed using the commercial Rapid ELISA Mouse mAb Isotyping kit (ThermoFisher). The kit determines the presence of the murine isotypes IgG1, IgG2a, IgG2b, IgG3, IgA, and IgM.

### Purification of mAbs

The purification of the mAbs took place in three phases: precipitation with PEG, molecular exclusion chromatography, and high-performance liquid exclusion and ion exchange liquid chromatography on an AKTA Purifier 10 (GE Healthcare). Precipitation with PEG was carried out by supplementing the culture supernatant with PEG 6000 to a final concentration of 4%. The suspension was kept under stirring for 3 h at room temperature and then subjected to centrifugation (1600 × *g*; 30 min at 4°C). The supernatant obtained after centrifugation was subjected to a second precipitation step with PEG 6000 6%, followed by centrifugation under the same conditions. The precipitate was dissolved in 15 mL of 50 mM Tris-HCl buffer solution, pH 8.0. The sample was fractionated by molecular exclusion chromatography (SEC) using the Superdex 200 High Load column (26 × 600 mm, 320 mL) with a flow rate of 3.0 mL / min, using 50 mM Tris-HCl, pH 8.0 as eluent. Collection volume was equal to 10 mL per tube. After selecting and pooling samples from the SEC, anion exchange chromatography was performed on a Poros HQ 10 × 100 mm column. Fractions were eluted under a flow of 5.0 mL / min with 50 mM Tris-HCl buffer solution, pH 8.0, in a gradient from 20 to 50% with saline. The fractions were collected with a volume of 4.0 mL per tube. The homogeneity of the samples obtained at each stage of the purification process was evaluated by denaturing electrophoresis on 12% polyacrylamide gel (SDS-PAGE) at a constant voltage of 200 volts for 45 min (Preserve & Flora Sherman 2011). To estimate the molecular weight (PM), the commercial standard Precision Plus ProteinTM Dual Color (Bio-Rad) was used. The proteins were developed with Coomassie Blue R350 dye solution and the result analyzed using the Image LabTM software, after image processing in the Gel DocTM XR + system (Bio-Rad).

### Antibody sequencing

The cells of each positive clone were collected by centrifugation (400 × *g* for 10 min at room temperature) and the RNA extracted with the commercial kit RNeasy Mini (Qiagen), following the protocol established by the manufacturer. For the Polymerase Chain Reaction via Reverse Transcriptase (RT-PCR), cDNA synthesis was performed from RNA using the commercial kit Super Script III First-Strand Synthesis System (INVITROGEN). Subsequently, PCR was performed with universal primers for murine VH and VL (42). RT-PCR was performed under the following conditions(42): initial denaturation 94°C / 5 min, denaturation 94°C / 2 min, annealing 48°C / 1 min, extension 72°C / 1 min and 30 sec. The cycles were repeated 35 times and the final extension was 72°C / 10 min for the VH chain and the same conditions for VL, but with the annealing temperature of 55°C / 1 min. The bands were visualized using 1.5% agarose gel (not shown). The sequencing of the selected mAbs was performed according with the protocol described in the commercial kit BigDye Terminator v3.1 (Life Technologies), and for that purpose the same primers described for PCR were used. The sequences were analyzed using the SeqMan program (DNAStar), and to identify the CDR1, 2 and 3, the gene sequences we used the IgBlast tool (IgBlast Tool - NCBI - NIH; https://www.ncbi.nlm.nih.gov/igblast/) through the Kabat database (http://www.bioinf.org.uk/abs/).

### Determination of affinity and dissociation constant by Plasmonic Surface Resonance (SPR)

The SPR experiments were performed using the BIACORE × system (GE Healtcare) equipped with a CM5 sensor chip. The ligands tested were the CC5 and DD11 mAbs, which were immobilized using amine coupling chemistry (Biacore × 202AD). The surfaces of the two flow cells were activated for 7 min with a 1: 1 mixture of 0.1 M NHS (N-hydroxysuccinimide) and 0.1 M EDC (3-(N, N-dimethylamino) propyl-N-ethylcarbodiimide) at a flow rate of 10 μL/min. The ligands were immobilized at 100 μg/mL in 10 mM sodium acetate, pH 5.0. Ester residues were deactivated with a 7 min injection of 1 M ethanolamine, pH 8.0. To collect kinetic binding data, the BSA-GlcNAc analyte was injected over the two flow cells in concentrations of 0.1 and 0.06 nM at a flow rate of 5 μL / min and at 25°C using HBS-EP (10 mM HEPES, 150 mM NaCl, 3 mM EDTA and 0.005% P20) pH 7.4. The data were adjusted by concentration in a simple model of interaction (1: 1) of the ligand and analyte using the option of global data analysis, which makes it possible to adjust all the graphs obtained simultaneously. All results were analyzed using the BiaEvaluation 4.1 software.

### ELISA for determination of antibody binding to intact cells

For this test, *C. neoformans, C. albicans, Giardia lamblia*, human lung cell line A549, *E. coli*, and *Staphylococcus aureus* were used. The fungal cells were washed in PBS three times and suspended at a density of 10^7^ cells/mL in poly-L-Lysine (5 μg/mL in PBS), and placed in 96-well polystyrene microplates for overnight incubation at 4°C (43). On the next day the plates were blocked with 5% PBS-BSA, for 1 h at 37°C, followed by incubation for 2 h at 37°C with the anti-chitooligomer mAbs at concentration ranges of 5 to 50 μg/mL. Subsequently, the systems were washed 3 times with PBS-Tween, and murine anti-IgM conjugated to peroxidase (1:5000 dilution) was added to the system, followed by incubation for 2h at 37°C. After washing, TMB was added and the plates were incubated for 30 min at 37°C. The reaction was stopped with 1M HCl 1, and readings were obtained at 450nm (43). Subsequently, the experiment was repeated, using serial dilutions (1:10) of the cell suspensions, which ranged from 10^2^ to 10^7^ cells/mL for fungi, and 10^4^ to 10^7^ cells / mL for the other cell types. The anti-chitooligomer mAb was tested at 25 μg/mL.

### Dot blot for determination of antibody binding to intact cells

Suspensions of *C. neoformans* and *C. albicans* were prepared at densities ranging from 10^2^ to 10^7^ cells/mL in 5 μg/mL poly-L-Lysine in PBS. These suspensions (10 μL) were loaded onto nitrocellulose membranes, and the steps of blocking and antibody binding were performed as in the ELISA. For reactivity determination, the membrane was cut into small pieces and deposited into the wells of 96-well plates, to which 50 μL of TMB were added. The systems were incubated for 30 min at 37°C. The solution (50 μL) was removed, transferred to a new plate and the development was carried out as described above (44).

### Anticryptococcal activity

Antimicrobial tests followed the EUCAST protocol adapted for yeast cells (17). *C. neoformans* cells were inoculated in RPMI 1640 buffered with 3 (n-morpholino) propanesulfonic acid (MOPS), pH 7, at 10^5^ cells/well in 96-well plates, in a final volume of 200 μL. The systems were supplemented with the mAbs (12.5; 6.25; 3.2; 1.6; 0.8; 0.4; 0.2; 0.1 μg/mL), amphotericin B (1 and 0.1 μg/mL). All substances were used alone or in combination with the mAbs. After 48 h of incubation at 37°C with shaking, the cells were homogenized with the aid of a pipette and subjected to reading at 592 nm (45). Synergistic activity between AmB alone and associated with the mAbs was determined on the basis of the calculation of the fractional inhibitory index (FII) and the synergism was categorized as follows: synergistic effect, FII < 1; additive effect, FII = 1 (46).

### Biofilm formation

The cell suspensions (*C. neoformans* H99) were prepared in minimal medium at 1 × 10^6^/mL and added (100 μL) to the wells of 96-well plates in the presence of the mAbs DD11 and CC5, all at 12.5 μg / mL. As a control of biofilm inhibition, amphotericin B was used at 1 μg/mL (45). Two independent systems were prepared; one of them were incubated for 48 h at 37°C, after that was washed to remove non-adherent cells and the fungal cells that remained attached to the wells were considered mature biofilms. While the other was not subjected to washes and the fungal cells were classified as biofilm formation. The first system after washes were incubated at 37°C for 24 h with antifungal drugs and mAbs; the second system the fungal cells was incubated with antifungal drugs and mAbs at 37°C for 48 h The metabolic activity of viable cells in the two assays was evaluated by the reduction of 2,3-bis-(2-methoxy-4-nitro-5-sulfophenyl)-2H-tetrazolium-5-carboxanilide (XTT), as determined spectrophotometrically at 492nm.

### Melanization

*C. neoformans* (1 × 10^6^ cells/mL) was cultured for 72 h in minimal medium supplemented with 1 mM L-DOPA in a 96-well plate (“U” bottom) in the presence of mAbs DD11 or CC5 at concentrations ranging from 0.2 to 25 μg/mL. Melanin production was determined densitometrically after digitalization of the images by the iBright FL1000 Invitrogen equipment (47).

### Murine infection

The influence of anti-chitooligomer antibodies on the survival of animals infected with lethal doses of *C. neoformans* was assessed using two different protocols. Mice (female Balb/C, 8 weeks old) (n = 7) were challenged i.p. with 10^5^ cells of *C. neoformans* H99 in PBS (100 µL/animal). After 2 h, the animals were treated i.p. with 100 μL of solutions of mAb DD11 at concentrations of 100, 250 or 500 μg/mL. As a control, the animals were treated with PBS only. The survival curve was monitored up to 90 days after infection. Alternatively, the mice were similarly infected, but treated after 2 h with 100 μL solutions of amphotericin B (2.5 mg/kg or 0.25 mg/kg)(48), mAb DD11 (85 μg/mL), or mAb DD11 (85 μg / mL) containing amphotericin B (0.25 mg / kg)(49). In all systems, the treatments were repeated twice at 10-day intervals. As controls, the animals were injected with 100 μL of PBS, AmB (2.5 mg / kg or 0.25 mg/kg) or MAb (85 μg / mL)(50). The survival curve was monitored up to 90 days after infection.

### Statistical analysis

All statistical analyses were performed using the GraphPad Prism software, version 5.00. Two-way ANOVA with Bonferroni post-test was used for individual comparison between groups, with a 95% confidence interval were used for all experiments. For the survival, curves, differences between groups were analyzed using the Gehan-Breslow-Wilcoxon test.

